# HiCzin: Normalizing metagenomic Hi-C data and detecting spurious contacts using zero-inflated negative binomial regression

**DOI:** 10.1101/2021.03.01.433489

**Authors:** Yuxuan Du, Sarah M. Laperriere, Jed Fuhrman, Fengzhu Sun

## Abstract

High-throughput chromosome conformation capture (Hi-C) has recently been applied to natural microbial communities and revealed great potential to study multiple genomes simultaneously. Several extraneous factors may influence chromosomal contacts rendering the normalization of Hi-C contact maps essential for downstream analyses. However, the current paucity of metagenomic Hi-C normalization methods and the ignorance for spurious inter-species contacts weaken the interpretability of the data. Here, we report on two types of biases in metagenomic Hi-C experiments: explicit biases and implicit biases, and introduce HiCzin, a parametric model to correct both types of biases and remove spurious inter-species contacts. We demonstrate that the normalized metagenomic Hi-C contact maps by HiCzin result in lower biases, higher capability to detect spurious contacts, and better performance in metagenomic contig clustering. The HiCzin software and Supplementary Material are available at https://github.com/dyxstat/HiCzin.

## 1 Introduction

High-throughput chromosome conformation capture (Hi-C) is a DNA proximity ligation approach with many applications in the investigation of genomic structures, DNA interactions, and even characterizing virus-host interactions from metagenomes [4, 14]. In Hi-C experiments, chimeric junctions are formed between pieces of DNAs in close proximity within cells and then subjected to paired-end sequencing generating millions of paired-end reads linking DNA fragments [14]. The number of reads connecting two DNA fragments is significantly related to the probability of contact between genomic loci in the three-dimensional structure at a fixed time point. Hi-C technique reveals the compartment property of the mammalian genomes [14], identifies topologically associated domains (TADs) [8], and reconstructs haplotypes [18].

Most recently, the Hi-C technique has been applied to the metagenomics domain (metagenomic Hi-C), and a series of Hi-C experiments have been conducted for microbial communities rather than a single species [7, 17]. Combined with the traditional shotgun sequencing, metagenomic Hi-C technique has displayed a powerful ability to probe virus–host interactions [4], simultaneously retrieve multiple genomes [6], deconvolute assembled contigs from whole genome shotgun (WGS) sequencing data into genome bins in both simulated and real microbial communities [2], and track horizontal gene transfer [20].

However, there exist strong experimental biases for the Hi-C interaction counts [21]; therefore, normalizing Hi-C data is essential to remove these biases. Though multiple strategies have been put forward [9, 10], most of these normalization methods aim to normalize Hi-C data derived from a single species, mainly human cells, and are not suitable to be applied on metagenomic Hi-C data from complex communities. This is mainly because potential factors of biases for metagenomic Hi-C data are different from those for Hi-C data within individual species. Additionally, it is not valid to theoretically assume all contigs should have equal visibility in metagenomic Hi-C data as the relative abundance levels of the different species can vary. Several relatively simple metagenomic Hi-C normalization methods have been developed. ProxiMeta [17] applied a normalization to the raw Hi-C counts by accounting for the estimated abundance of the contigs, and further took the number of restriction sites on the contigs into consideration [19]. As a proprietary metagenomic genome binning platform without open-source pipeline, ProxiMeta did not clarify the normalization algorithms in detail. Beitel et al. [3] divided raw interaction counts by the product of the length of two contigs. MetaTOR [2] normalized raw counts by the geometric mean of the contigs’ coverage. Metaphase [6] and Bin3C [7] divided raw Hi-C counts by the product of the number of restriction sites and Bin3C used the Knight-Ruiz algorithm [12] to construct a general doubly stochastic matrix after the first step correction. We will show that these normalization methods are not effective in removing all biases. Additionally, the biases of spurious inter-species contacts are ignored for metagenomic Hi-C data by all these normalization methods, considerably weakening the interpretability of the Hi-C data [19].

Here we first comprehensively discuss potential experimental biases for metagenomic Hi-C data, and then propose HiCzin, a method to normalize metagenomic Hi-C data based on the zero-inflated negative binomial regression frameworks [22]. We also develop a hybrid statistical method to detect spurious inter-species contacts. We show that the normalized metagenomic Hi-C contact maps by HiCzin lead to lower biases, higher ability to detect spurious contacts, and better performance in metagenomic contig clustering on the published metagenomic Hi-C dataset.

## 2 Result

### 2.1 Source of biases in metagenomic Hi-C experiments

In addition to chromosomal contacts of interest, several other factors unrelated to chromosomal contacts can also influence the number of Hi-C interactions between contigs [21]. We refer to such factors as biases. We report on two kinds of biases with substantial influences on metagenomic Hi-C contact maps: explicit biases and implicit biases. Explicit biases include three potential factors: i) the number of enzymatic restriction sites on contigs, ii) contig length, and iii) contig coverage [3, 6, 17], all of which can be observed. Implicit biases include unobserved interactions and spurious inter-species contacts. Unobserved interactions are chimerical DNA fragments that are missed due to the factors such as the mappability of contigs and in vivo constraints on accessibility. Spurious inter-species contacts arise from the ligation of DNA fragments between closely related species [19]. As implicit biases are unobservable, it is challenging to detect and correct implicit biases.

### 2.2 Analyses of experimental biases in synthetic metagenomic yeast samples

We analyzed metagenomic yeast (M-Y) samples, consisting of 16 yeast strains (BioProject : PRJNA245328) [6]. After processing the raw WGS and Hi-C reads, we generated raw Hi-C contact maps for 6,196 assembled contigs (Supplementary Material). Reference genomes of these 16 yeast strains were downloaded (Supplementary Material: Table S1). To determine the true species identity of the assembled contigs, contigs were aligned to reference genomes at the species level by BLASTn [1] (Supplementary Material). Thirty-seven contigs (0.6%) could not be aligned to reference genomes and were not considered in the following analyses (Supplementary Material: Figure S1).

According to the alignment results to the reference genomes, we refer to contig pairs from the same species and different species as intra-species pairs and inter-species pairs, respectively. Interaction counts of intra-species pairs and inter-species pairs are defined as valid contacts and spurious contacts, respectively. In particular, we denote zero contacts if no interaction was observed between intra-species pairs; hence the intra-species contacts, corresponding to intra-species pairs, are composed of valid contacts and zero contacts. Valid contacts imply a high probability of contig pair’s belonging to the same genome, while spurious contacts confound the interpretation of the Hi-C data.

Raw interaction counts were enriched between pairs of contigs with a high number of restriction sites, long contigs, and/or contigs with high coverage (Figure 1), which can be explained by the following reasons. Longer contigs may have higher ligation efficiencies with other contigs than shorter contigs, more restriction sites are likely to increase the probability of enzymatic cuts within DNA fragments, and higher coverages, representing higher concentration of contigs, can result in more Hi-C interactions between contigs. The Pearson correlation coefficients between raw valid contacts and the product of the number of restriction sites, the length and the coverage for each pair of contigs were 0.429, 0.400, and 0.184, respectively, demonstrating that these three factors were indeed highly correlated with valid contacts.

**Fig. 1.**
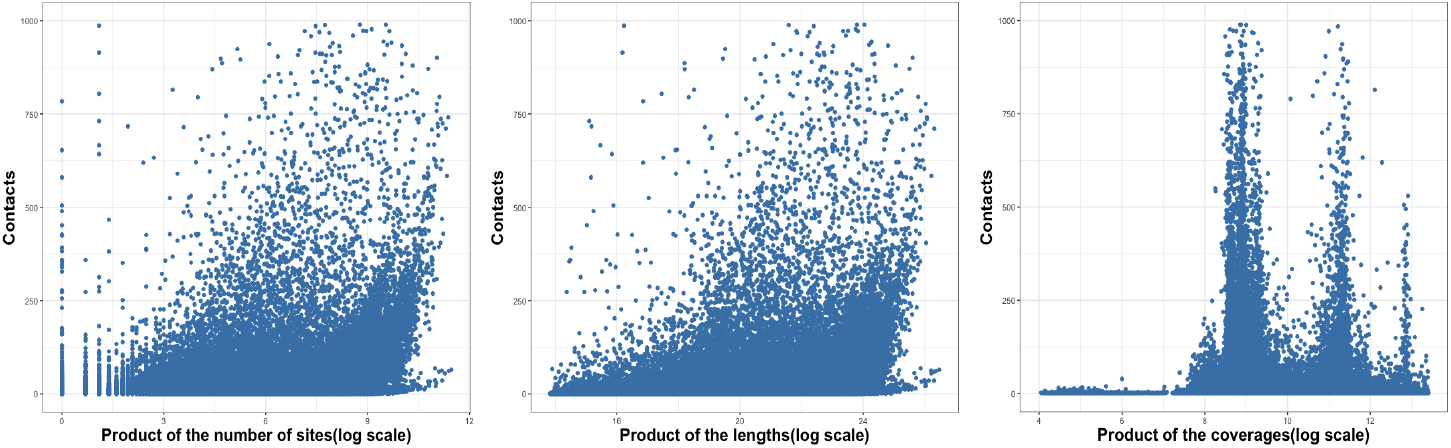
Relationship between raw interaction counts and the product of the number of restriction sites, length, and coverage between contig pairs.

As for implicit biases, one remarkable phenomenon for intra-species contacts was the presence of excess zeros, which means zero contacts account for a large proportion within intra-species contacts. The number of valid contacts (i.e., nonzero intra-species contacts) only made up 14.9% within all intra-species contacts, suggesting the potential existence of unobserved interactions with high probability due to the experimental noise. The number of spurious contacts made up 25.5% of all nonzero contacts, which could not be neglected for the M-Y samples.

### 2.3 Normalization methods in the publicly available metagenomic Hi-C analysis pipelines

Because of the existence of aforementioned experimental biases, it is necessary to normalize the raw Hi-C contacts before downstream analysis, such as clustering and tracking virus-host interactions. Most of the current available pipelines divided the raw Hi-C interactions by the product of one factor of explicit biases to normalize raw Hi-C contacts, which we refer to as naive normalization methods [2, 3, 17]. These naive normalization methods only corrected part of explicit biases, and the unnormalized factors of explicit biases might still be highly correlated with Hi-C contact maps. As for the two-stage normalization method in Bin3C [7], equal visibility for all regions is a basic theoretical assumption for utilizing the matrix balancing algorithm they use to recover normalized Hi-C matrices [10], yet this assumption is not satisfied for metagenomic assembled contigs with huge differences in length and abundance. Moreover, all these normalization methods ignored the influence of implicit biases and did not attempt to detect and remove the spurious inter-species contacts. Therefore, it is imperative to develop new normalization methods to overcome these shortcomings.

### 2.4 Removing explicit biases and spurious contacts using a zero-inflated negative binomial regression framework

The Poisson and negative binomial regression models are widely used in fitting count data and have been successfully employed in fitting Hi-C interactions of human cells [9]. Therefore, there is potential to apply frameworks based on Poisson or negative binomial regression to normalize metagenomic Hi-C data. Here we model the population of the intra-species contacts using the negative binomial distribution rather than the Poisson distribution because Hi-C data are always over-dispersed [9]. In the classical negative binomial regression model, we can fit the model given sample data of the intra-species contacts by regarding factors of biases and intra-species contacts as predictor variables and the response variable, respectively. Then, the residuals of this conventional model serve as normalized contacts.

However, some underlying interactions may not be observed in Hi-C experiments due to the limited quantity of Hi-C reads and problems in mapping Hi-C reads to the contigs. Ignoring such influences may lead to serious biased estimation and prediction. Additionally, although classical negative binomial models can capture the property of over-dispersion, they are not sufficient for modeling the excess zeros observed in Hi-C contact maps. To solve these problems, we developed HiCzin, a novel metagenomic Hi-C normalization method based on zero-inflated negative binomial regression frameworks [22], combining the counting distribution of the intra-species contacts with a mass distribution of unobserved contacts. The residues of the counting part serve as normalized contacts (see Methods).

Compared with raw valid contacts, the average value of raw spurious contacts was smaller (Figure 2a), while the average number of restriction sites, length, and coverage of contigs were significantly larger (Figure 2b,c,d). These evidences indicated that spurious inter-species contacts were more likely to be generated for longer contigs with more restriction sites and higher abundances. Therefore, we expect that the magnitude of the normalized spurious contacts by the factors of explicit biases to be significantly smaller than that of the normalized valid contacts. Thus, a basic idea is to discard the normalized contacts whose values are less than a selected threshold as spurious contacts [19]. However, determining the threshold is extremely challenging. Based on our HiCzin normalization model, we develop a hybrid statistical method to detect spurious contacts and determine thresholds (see Methods).

**Fig. 2.**
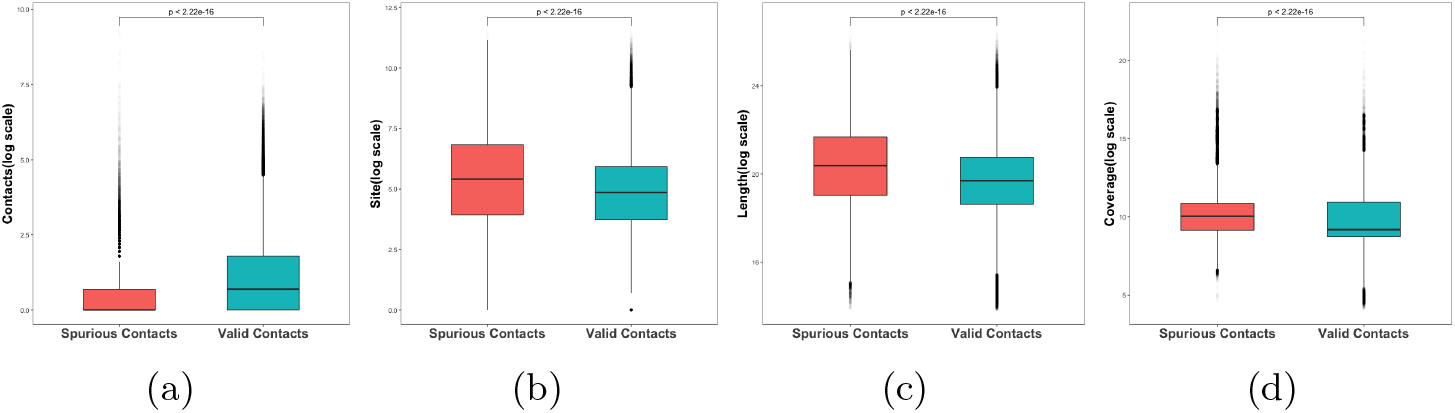
Comparison of (a) the raw counts of spurious contacts and valid contacts, (b) the number of restriction sites of spurious contacts and valid contacts, (c) the length of spurious contacts and valid contacts, (d) the coverage of spurious contacts and valid contacts.

### 2.5 Applying the HiCzin model to the M-Y samples

To fit the HiCzin model, samples of the intra-species contacts were generated using TAXAassign (https://github.com/umerijaz/TAXAassign), which assigned 3,441 (55.5%) contigs to the known reference genomes in the NCBI nt database (see Methods). These 3,441 contigs were assigned to 10 species by TAXAassign (Supplementary Material: Figure S2). We compared the taxonomy assignment results by TAXAassign with the corresponding true species identities obtained by BLASTn. Only 21 labels were different, indicating the high precision of taxonomy assignments at the species level by TAXAassign. Then, taking advantage of these labeled contigs, we generated a relationship of intra-species pairs by pairwise combining contigs from the same species, and corresponding contacts were obtained as sample data to fit the HiCzin model. A total of 1,492,856 samples of the intra-species contacts were generated.

All sample data were then utilized to fit the HiCzin model. We compared our model with naive normalization methods, the two-stage normalization method in Bin3C and the classical negative binomial regression model. To simplify the notation, we denote naive normalization methods by site, length, and coverage as Naive Site, Naive Length, and Naive Coverage, and denote the two-stage normalization method in Bin3C and the classical negative binomial regression model as Bin3C Norm and Naive NB.

We first calculated the Pearson correlation coefficients between normalized valid contacts and the product of each of the three factors of explicit biases to gauge the bias effects (Table 1). The Naive Site and Naive Length approaches increased the Pearson correlation coefficients between valid contacts and the product of the coverage from 0.184 to 0.559 and 0.694; the Naive Coverage approach increased the correlation coefficient between valid contacts and the product of the site from 0.429 to 0.515 and increased the correlation coefficient between valid contacts and the product of the length from 0.400 to 0.481. These results proved that the naive normalization methods only corrected part of explicit biases, and the unnormalized factors of explicit biases showed even higher correlation with Hi-C contact maps. In contrast, the two-stage normalization method in Bin3C decreased all three correlation coefficients to 0.024, 0.025, 0.011, indicating that the matrix balancing algorithm can assist in correcting explicit biases to some extent. These three correlation coefficients were decreased to 0.023, 0.024, 0.154 using Naive NB, and further decreased to 2 ×10^−4^, 0.002, 0.069 using HiCzin. Therefore, HiCzin achieved better performance than all other normalization methods in removing explicit biases.

**Table 1.**
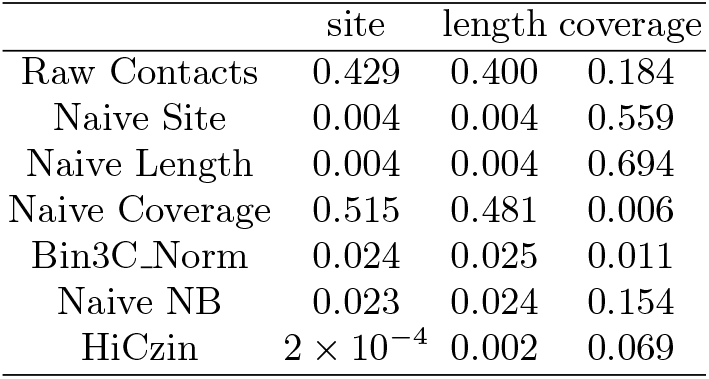
Pearson correlation coefficients (absolute value) between normalized valid contacts and the product of each of the three factors of explicit biases.

The other objective of metagenomic Hi-C normalization is to identify valid contacts from all observed contacts. Although raw values of spurious contacts were significantly smaller than those of valid contacts, the distribution of spurious contacts mixed with the distribution of valid contacts (Figure 3a), making it challenging to separate spurious contacts from valid contacts. After normalization, the distribution of normalized spurious contacts deviated considerably to the left from the distribution of normalized valid contacts (Figure 3b), facilitating the distinction from spurious contacts to valid contacts.

**Fig. 3.**
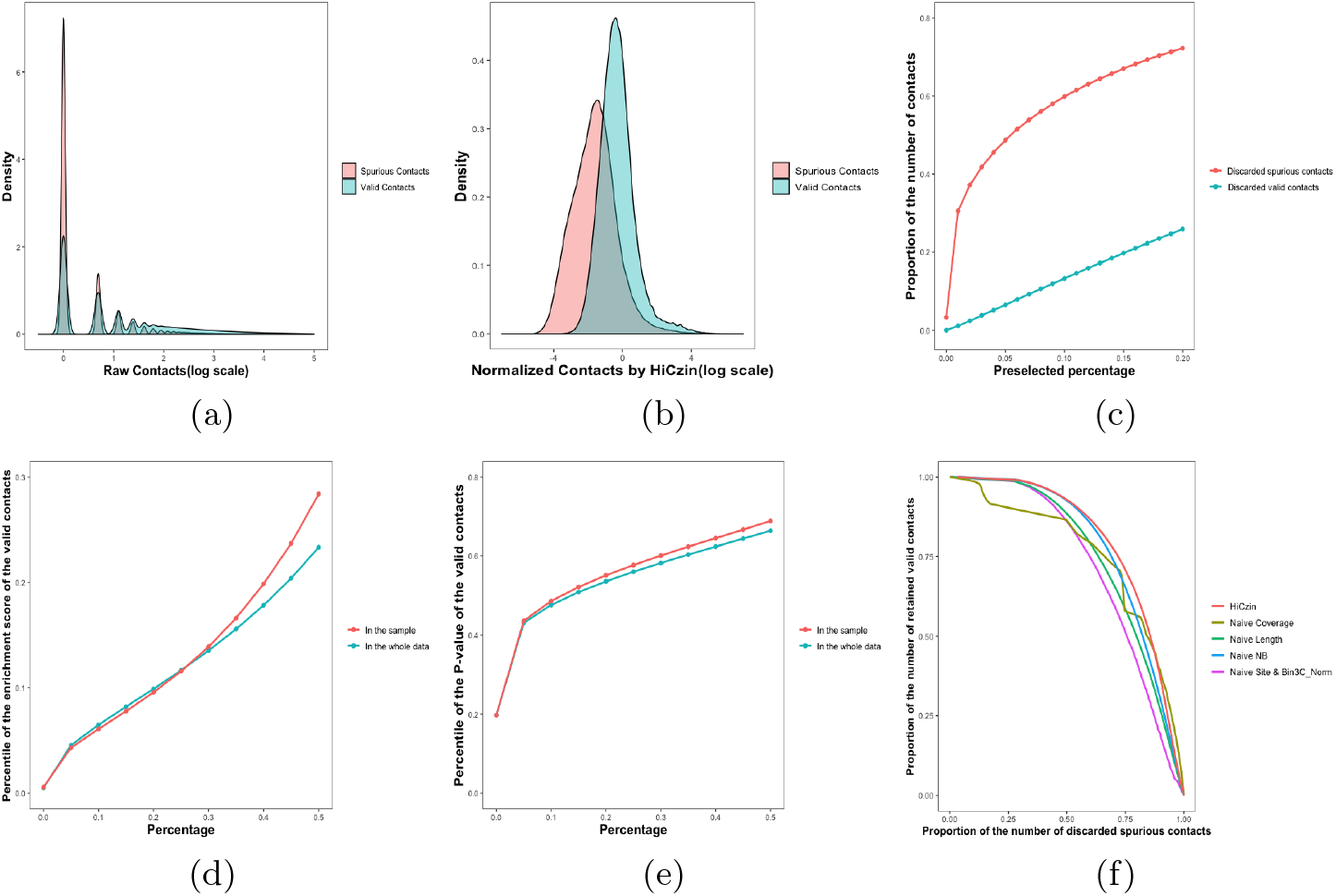
(a) Comparison of the distribution of raw valid contacts and raw spurious contacts. (b) Comparison of the distribution of normalized valid contacts and normalized spurious contacts by HiCzin. (c) The proportions of discarded valid contacts and discarded spurious contacts. (d) The percentiles of the enrichment score of the valid contacts in the sample and in the whole data. (e) The percentiles of the *p*-value of the valid contacts in the sample and in the whole data. (f) The discard-retain curve using all sample data.

Therefore, we adopted our hybrid statistical approach based on the HiCzin model to detecting and then discarding spurious contacts (see Methods). The main procedure of our approach is to select thresholds of the enrichment score and the *p*-value, respectively, and any contacts whose enrichment score or *p*-value are below the thresholds would be identified as spurious contacts. A percentage reflecting the acceptable fraction of losses of the valid contacts was preselected and thresholds were determined such that less than the preselected percentage of valid contacts in sample data were incorrectly identified as spurious contacts. Noticeably, both thresholds increased with the preselected percentage, and larger thresholds could detect more spurious contacts while incorrectly identifying a higher number of valid contacts. Though there existed a ‘trade-off’, the proportion of discarded spurious contacts increased much faster than that of discarded valid contacts (Figure 3c), indicating that we could remove a large fraction of spurious contacts while keeping most of the valid contacts. For instance, if we set the preselected percentage as default 10%, which means that we could withstand the losses of around 10% of valid contacts, about 60% of spurious contacts were detected while only 13% of valid contacts were incorrectly removed. In addition, the percentiles of the enrichment score and *p*-value of the valid contacts in our sample data were indeed close to those in the whole data (Figure 3d,e), ensuring that our method to determine the thresholds by utilizing the sample data could make the proportion of incorrectly discarded valid contacts in the whole data under control. These results supported the feasibility of our spurious contact detection method.

According to our basic idea of spurious contact detection, naive normalization methods and the Naive NB method could also be used to detect spurious contacts by regarding normalized contacts less than certain thresholds as spurious contacts. We employed the same technique proposed in our hybrid statistical spurious contact detection method to determine thresholds for other normalization methods. For the two-stage normalization method in Bin3C, as the matrix balancing algorithm in the second step may amplify the influence of certain spurious contacts, it is better to remove the noise of spurious contacts after the first step correction. Since the first stage of Bin3C Norm is equivalent to the Naïve Site approach, the spurious contact detection result of Bin3C Norm is the same as that of the Naive Site approach. To evaluate the capability of normalization methods to detect spurious contacts while retaining the valid contacts, we design the discard-retain curve (DR curve). In the graph of a DR curve, the x-axis is the proportion of discarded spurious contacts among all spurious contacts in the whole data, and the y-axis represents the proportion of retained valid contacts within all valid contacts in the whole data. We denote the area under the DR curve as AUDRC. Larger AUDRC indicates that the normalization method can retain more valid contacts while discard more spurious contacts. Therefore, we plotted the DR curve to evaluate the performance of different normalization methods (Figure 3f). AUDRC was subsequently calculated for each of the normalization methods (Table 2), and our HiCzin model achieved the best result with respect to AUDRC.

**Table 2.**
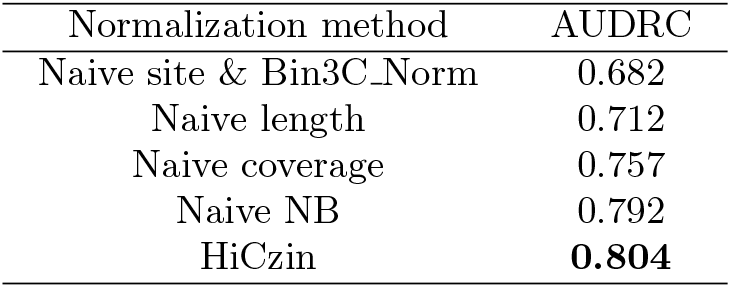
Area under the discard-retain curve for different normalization methods. Higher AUDRC score indicates better performance in spurious contact detection.

### 2.6 Evaluation of the impact of the number of labeled contigs and detected species on the HiCzin model

The above results showed that the HiCzin model achieved outstanding performance in Hi-C normalization and spurious contact detection by utilizing all sample data to fit the normalization model. Around half of the contigs corresponding to 10 out of 13 species were labeled by TAXAassign, providing us with enough samples to fit the model. However, in real situations, we sometimes can only label a small number of contigs at the species level and detect a low quantity of species.

To explore how the HiCzin model performs with only a small number of labeled contigs and detected species, we consequently removed labeled contigs belonging to certain species step by step. Specifically, we first removed three species (S.mikatae, P. pastoris, L. kluyveri) to which only a small number of contigs (*<*20) were assigned. The remaining seven species were sorted in descending order by the number of contigs assigned to these seven species. Then, we removed the labeled contigs of one species at a time in the above species’ order until fewer than 10% of all contigs remained. In this way, we could simulate both situations where some species are unknown, and only a small number of contigs can be labeled (Supplementary Material: Table S2).

Although the performance of normalization and spurious contact detection became slightly worse as the number of labeled contigs and detected species decreased, the results were still better than the naive normalization methods and the two-stage normalization methods in Bin3C for the different sample sizes (Supplementary Material: Table S3). The HiCzin model also obtained better performance in the spurious contact detection (Supplementary Material: Figure S3). Therefore, the HiCzin model can achieve good results even when the number of labeled contigs and detected species are relatively low.

### 2.7 Generalizing the HiCzin model

Our HiCzin model can be generalized to consider different independent variables and do normalization without labeled contigs (see Methods). Here, we explore three significant scenarios.

#### HiCzin LC

In some real situations, the specific enzymes utilized in Hi-C experiments are unknown; thus only the length and the coverage of contigs can serve as independent variables.

#### HiCzin GC

GC-content is sometimes considered as one source of biases in Hi-C experiments [9, 21]. Therefore, we explored the influence of adding GC-content as a new predictor variable to our HiCzin model, though we did not observe a strong correlation between raw valid contacts and GC-content (Pearson correlation coefficient: 0.032) for the Hi-C contact maps of the synthetic M-Y samples.

#### Unlabeled HiCzin

In the real application of HiCzin, some extreme difficulties may be encountered. For example, there may not be enough computational resources to run TAXAassign or an extremely small number of contigs can be labeled. To solve these problems, a HiCzin normalization mode without labeled contigs (Unlabeled HiCzin) is designed.

We applied these three generalized HiCzin models on the M-Y samples (Supplementary Material: Table S4). For the HiCzin LC and HiCzin GC, the Pearson correlation coefficients between normalized contact counts and the three factors increased compared to those of the HiCzin in Table 1, though the AUDRC was slightly higher than that of the HiCzin in Table 2. For the unlabeled HiCzin, detecting spurious contacts was tough as it was challenging to determine thresholds without specific samples of the intra-species contacts. Although the normalization results were worse than those of the HiCzin model using labeled contigs, unlabeled mode of HiCzin still performed better than naive normalization methods and it is more applicable and requires fewer computational resources than the HiCzin model using labeled contigs.

### 2.8 Clustering of contigs by the Louvain algortihm

The Louvain algorithm has been widely employed to cluster contigs based on metagenomic Hi-C data [2, 16]. We applied this algorithm to the Hi-C data normalized by different methods. We set the preselected percentage of maximum incorrectly identified valid contacts in sample data as 10% for all HiCzin models and regarded groups above 500 kbp as effective bins to evaluate the clustering performance.

As shown in Table 3, the original HiCzin model achieved the best clustering performance by the Louvian algorithm. Although the matrix balancing algorithm in the second stage could improve the clustering quality, Bin3C Norm grouped much fewer contigs compared to Naive Site. The Naive NB approach also grouped a relatively small number of contigs. The performance of both correcting biases and clustering by HiCzin GC was worse than that of the original HiCzin model; hence it is not necessary to consider the GC-content in the regression process. One potential explanation for the poor performance of HiCzin GC is that the genomes in the community have similar GC-content and the Hi-C contact maps are not dependent on GC-content. The clustering results of the HiCzin LC and the unlabeled HiCzin were significantly better than all naive normalization methods and the two-stage normalization method in Bin3C, indicating that our normalization model still performed well when the restriction enzymes of Hi-C experiments or the labels of any contigs were unknown. These results ensure that the HiCzin models are widely applicable with excellent normalization effects under different circumstances.

**Table 3.**
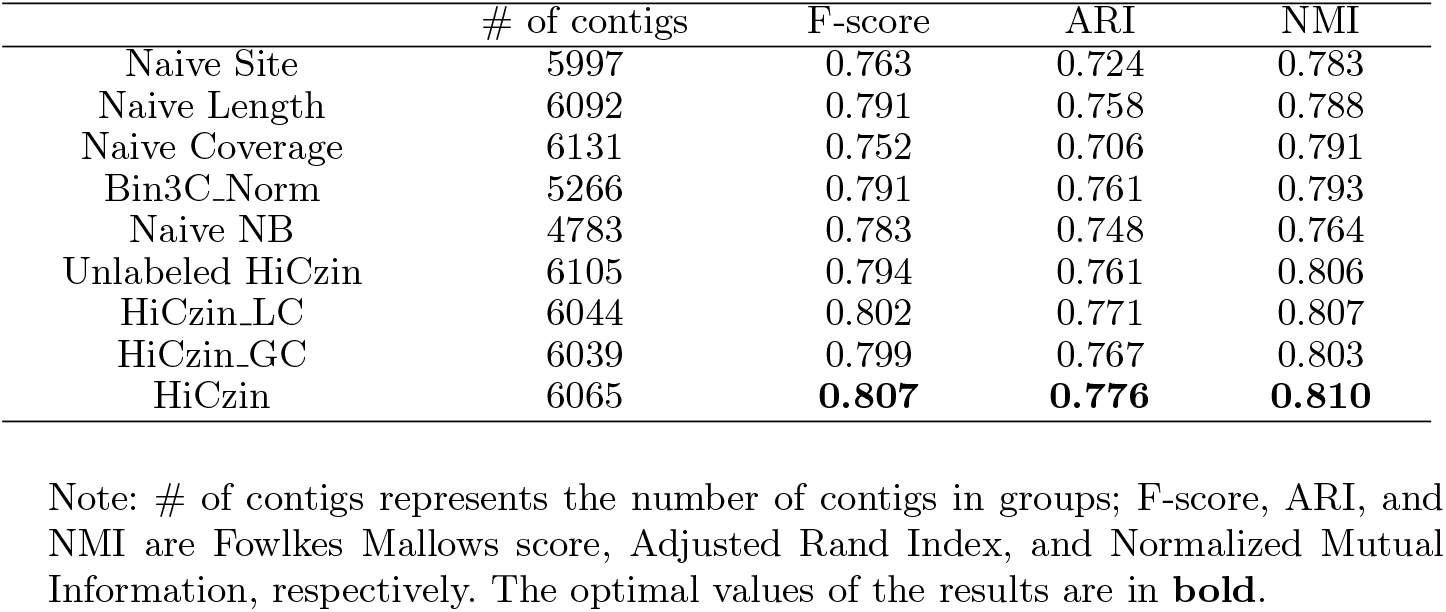
Comparison of the clustering results of contigs using the Louvain Algorithm.

## 3 Methods

### 3.1 Framework of applying HiCzin to metagenomic Hi-C experiments

The workflow of HiCzin utilized in the metagenomic Hi-C analysis is shown in Figure 4. In metagenomic Hi-C experiments, short reads are obtained by shotgun sequencing from microbial communities. At the same time, metagenomic Hi-C sequencing reads are generated from the same sample. Contigs are assembled from the shotgun short reads and Hi-C reads are mapped to the assembled contigs to construct raw contact maps consisting of the number of Hi-C reads mapped to contig pairs. Then, HiCzin is employed to normalize raw contact maps and discard spurious contacts. Finally, downstream analysis can be conducted on the basis of normalized contact maps by HiCzin.

**Fig. 4.**
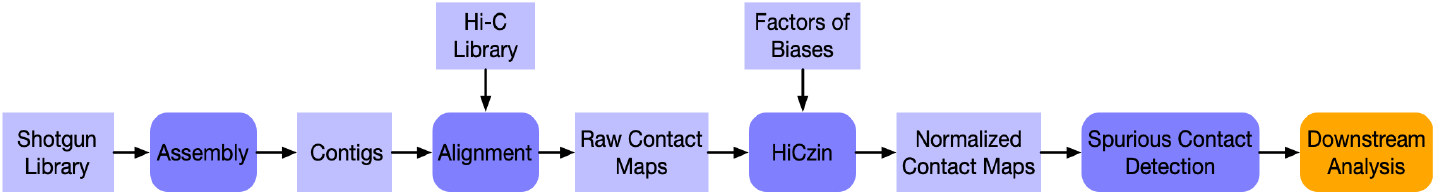
Workflow of HiCzin utilized in metagenomic Hi-C analysis

### 3.2 Calculating the coverage of assembled contigs

The coverage of contigs was computed using MetaBAT [11] v2.12.5 script: ‘jgi_summarize_bam_contig_depths’.

### 3.3 Applying TAXAassign to generate sample data of the intra-species contacts

The taxonomic assignment of contigs was resolved using NCBI’s Taxonomy and its nt database by TAXAassign(v0.4) with parameters ‘-p -c 20 -r 10 -m 98 -q 98 -t 95 -a “60,70,80,95,95,98” -f’. Assignment results with ‘unclassified’ at the species level were discarded, and only deterministic results of taxonomic assignment at the species level were kept. Intra-species pairs were subsequently generated by pairwise combining contigs assigned to the same species, and corresponding contacts were treated as samples to fit the HiCzin model.

### 3.4 Normalization via the HiCzin model

Based on the zero-inflated generalized linear mixed framework [13], the HiCzin is a two-component mixture model combining a mass point at zero with a count distribution. Specifically, within the intra-species contacts, zero contacts may come from two sources: the count distribution, showing that these zeros are observations of the population of the intra-species contacts and no interactions happened, or the zero mass points, indicating that Hi-C interactions happened, but the observations of the interactions were lost due to certain kinds of experimental noise.

Formally, denote the population of the intra-species contacts as a random variable *Y*. The basic assumption of the HiCzin model is that *Y* follows the negative binomial distribution. Let *π*_*ij*_ denote the probability of unobserved contacts and *Z*_*ij*_ denote a zero-inflated random variable of the intra-species contacts between the *i*th contig and the *j*th contig. Then the random variable *Z*_*ij*_ is given by

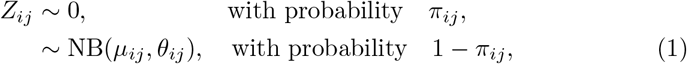

where NB(*µ*_*ij*_, *θ*_*ij*_) is negative binomial distribution with mean *µ*_*ij*_ and shape parameter *θ*_*ij*_.

Therefore, the zero-inflated density of *Z*_*ij*_ is the result of mixing a negative binomial distribution and a degenerate distribution at zero as

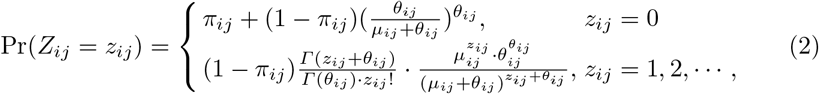

where *Γ* (·) is the gamma function. The random variable *Z*_*ij*_ will be degenerated to negative binomial distribution when *π*_*ij*_ = 0.

We assume that the parameters *µ*_*ij*_ and *π*_*ij*_ depend on the three factors of explicit biases while *θ*_*ij*_ is an independent parameter as a constant parameter *θ* in our model. Define *s*_*k*_, *l*_*k*_, and *c*_*k*_ as the number of restriction sites, the length and the coverage of the *k*th contig, respectively. As Lord et al. [15] suggested, link functions in generalized linear models are used to model the dependence of parameters *µ*_*ij*_ and *π*_*ij*_ on the three factors of explicit biases. To be specific, we propose that *µ*_*ij*_ is related to three factors by the logarithmic link, i.e.,

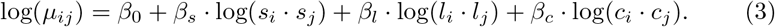

We also propose that *π*_*ij*_ is releted to three factors by the logistic link, i.e.,

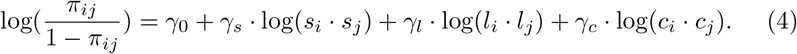

Let 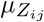 denote the mean of random variable *Z*_*ij*_. Then, the corresponding regression equation for 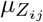 is

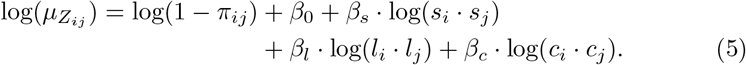

The overall model parameters *β* = (*β*_0_, *β*_*s*_, *β*_*l*_, *β*_*c*_), *γ* = (*γ*_0_, *γ*_*s*_, *γ*_*l*_, *γ*_*c*_) and additional dispersion parameter *θ* can be estimated by maximum likelihood (ML) using the latest R package ‘glmmTMB’ [5].

Finally, the residuals of the counting part are the normalized metagenomic Hi-C contacts, i.e.,

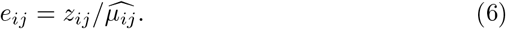

Hence, given sample data of the intra-species contacts, our HiCzin model can integrate all three factors of explicit biases. The influence of unobserved interactions is also taken into account simultaneously to ‘unbiased’ the estimation and prediction.

### 3.5 Spurious contact detection by a hybrid statistical method based on HiCzin

From the HiCzin model, the intra-species contact *Y*_*ij*_ follows the negative binomial distribution with mean 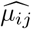 and shape 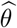. Given any contig pairs with nonzero contacts, we denote the value of the observed raw contacts as *O*_*ij*_ and the expected contacts under condition that the two contigs come from the same species as *E*_*ij*_, where 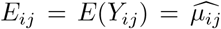. We define the enrichment score as *S*_*ij*_ = log(*O*_*ij*_*/E*_*ij*_).

Under our statistical framework, we also design a hypothesis test to detect spurious contacts. Since observations of Hi-C interactions need to be protected, the null hypothesis of the test is that *O*_*ij*_ belongs to the intra-species contacts while the alternative hypothesis is that *O*_*ij*_ belongs to the spurious inter-species contacts. We directly regard *O*_*ij*_ as the test statistic and *O*_*ij*_ ∼ *Y*_*ij*_ under null hypothesis. We choose one-tailed test and calculate the *p*-value of *O*_*ij*_ as

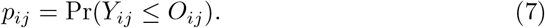

Then, we develop a hybrid statistical method to detect spurious contacts. We choose a threshold *t* for the enrichment score and a significance level *α* for the hypothesis test. Contacts of contig pairs whose enrichment score is less than *t* or *p*-value is less than *α* will be regarded as spurious contacts and then discarded.

To determine the threshold and the significance level, we assume that the percentiles of the enrichment score and the *p*-value of the valid contacts in our sample data are similar to those in the whole data and preselect a percentage (default 10%) reflecting the acceptable fraction of losses of the valid contacts. Taking advantage of our generated sample data of the intra-species contact, we can determine the threshold *t* and significance level *α* such that less than the preselected percentage of valid contacts in sample data are incorrectly identified as spurious contacts for both methods, respectively. Based on our assumption, we suppose that around the same percentage of valid contacts in the whole data might be mistakenly discarded. Therefore, thresholds can be strictly restricted to detect most of spurious contacts while avoid incorrectly identifying a large proportion of valid contacts in the whole data.

### 3.6 Generalizing the HiCzin by selecting different independent variables

Let 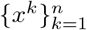 denote the set of factors. Then, we modify the regression equation in (5) as

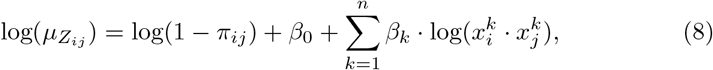

where *π*_*ij*_ in (4) is modified as

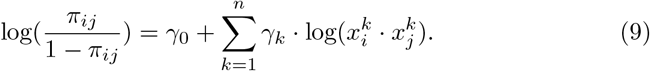

Then, ‘glmmTMB’ [5] package is employed to estimate 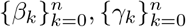, and *θ*, and residuals of the counting part are considered as normalized contacts.

### 3.7 HiCzin normalization without labeled contigs

As samples of the intra-species contacts cannot be obtained in some scenarios, we just regard all nonzero raw contacts as our sample data. Although these samples contain both valid contacts and spurious contacts, the number of valid contacts is supposed to be much larger than that of spurious contact and thus we suppose that spurious contacts will not result in significant biases in parameter estimation. Moreover, as we don’t have zero contacts to fit the zero-inflated part, one option to solve this problem is to set *π*_*ij*_ as a constant parameter, i.e.,

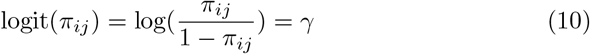

Unknown parameters are estimated by maximum likelihood using ‘glmmTMB’ [5] package and residuals are considered as normalized contacts.

## 4 Conclusions and Discussions

We put forward two types of experimental biases for metagenomic Hi-C data. Explicit biases include the number of restriction sites, contig length, and contig coverage and implicit biases include unobserved interactions and spurious inter-species contacts. Both types of biases could be obviously observed in the metagenomic yeast samples. Naive normalization methods could only correct part of explicit biases, and the unnormalized factors of explicit biases showed even higher correlation with Hi-C contact maps. Based on the basic assumption that the population of the intra-species contacts follows the negative binomial distribution, we have presented HiCzin, a parametric model applying zero-inflated negative binomial regression framework to normalize metagenomic Hi-C data, and have introduced a hybrid statistical method to detect and remove the spurious inter-species contacts. The HiCzin model takes the impact of unobserved interactions into account. We have shown that normalized metagenomic Hi-C contact maps by HiCzin lead to lower biases, higher ability to detect spurious contacts, and better metagenomic contig clustering performance, compared with all naive methods and two-stage normalization method in Bin3C. In case that the specific enzymes utilized in Hi-C experiments are unknown or there are not enough computational resources to run TAXAassign, we come up with the generalized HiCzin by only selecting the length and the coverage of contigs as predictor variables, and a HiCzin mode without labeled contigs. We have shown that these two models also performed well in normalization, spurious contact detection and metagenomic contig clustering. Although we can remove a large fraction of spurious contacts by our hybrid statistical approach, it is inevitable to lose a small quantity of useful valid contacts. Directly modeling the spurious contacts may separate the spurious contacts from valid contacts even better. As the Hi-C technique will be increasingly utilized upon the metagenomics domain in the near future, we expect that the normalization model we propose here can facilitate the downstream analysis, and improve results in retrieving metagenome-assembled genomes, identifying virus-host interactions, tracking horizontal gene transfer and all other areas making use of metagenomic Hi-C data.

