## Supplementary Material for "HiCzin: Normalizing metagenomic Hi-C data and detecting spurious contacts using zero-inflated negative binomial regression"

[1] Department of Quantitative and Computational Biology, University of Southern California, USA
{yuxuandu,fsun}@usc.edu
[2] Department of Biological Sciences, University of Southern California, USA

### Materials

Metagenomic yeast (M-Y) samples are synthetic microbial samples with 16 yeast strains and 13 yeast species. Raw WGS and Hi-C libraries were constructed by Metaphase [2]. WGS dataset contains 85.7 million read pairs (101 bp per read) and Hi-C dataset contains 81 million read pairs (100 bp per read).

### Initial processing

We applied a standard cleaning pipeline on both WGS and Hi-C datasets using bbduk from the BBTools suite(v37.25) [3]. Adaptor sequences were removed by bbduk with parameter 'ktrim=r k=23 mink=11 hdist=1 minlen=50 tpe tbo' and reads were quality-trimmed using bbduk with parameter 'trimq=10 qtrim=r ftm=5 minlen=50'. We recommend using softwares(e.g. DeconSeq [7]) to remove noise from human if necessary [8]. Then, the first 10 nucleotides of each read were trimmed by bbduk with parameter 'ftl=10'.

### Shotgun assembly and Hi-C read alignment

For the shotgun dataset, metagenome assembly was produced by MEGAHIT [4] with parameters '-min-contig-len 300 -k-min 21 -k-max 141 -k-step 12 -merge-level 20,0.95' and contigs shorter than 1 kb were discarded.

For the Hi-C dataset, only paired reads were kept for the downstream analysis. All PCR optical and tile-edge duplicates for Hi-C paired-end reads were removed by 'clumpify.sh' from BBTools suite [3] with default parameters. Processed Hi-C paired-end reads were mapped to assembled contigs using BWA-MEM [5] with parameters '-5SP'. Then, samtools [6] with parameters 'view -F 0x904' were applied on the resulting BAM files to remove unmapped reads (0x4) and supplementary (0x800) and secondary (0x100) alignments. Alignments with low quality ( <30 nucleotide match length or mapping score <30) were also filtered out.

By this means, 4,700,202 read pairs were mapped to different contigs for the synthetic M-Y samples.

### Contact map generation

As contact map reflects the proximity distance within contigs, only pairs of reads aligned on different contigs were kept so as to generate the contact map. Raw contig–contig interactions were aggregated as contacts by counting the number of alignments linking two contigs. Contigs that no Hi-C reads were aligned to were discarded.

### Annotating contigs by reference genomes for the M-Y samples

In order to explore experimental biases, the reference genomes of 16 yeast stains were downloaded (Table S1). As analysis was performed at the species level, the genomes of four strains(FY, CEN.PK, RM11-1A and SK1) from the same species (S. cerevisiae) were combined into one reference genome. Then, all contigs were aligned to those 13 reference genomes of all known species by BLASTn [1] with parameters: '-perc_identity 95 -evalue 1e-30 -word_size 50'. Hence, the true species that assembly contigs came from were determined if there existed any alignment of the contigs to the species' reference genome; the placement of the alignment was ignored [2].

### Evaluation criteria of clustering

**Fowlkes-Mallows scores:** The Fowlkes-Mallows score (FM) is defined as the geometric mean of the pairwise precision and recall, i.e,

$$\text{FM} = \sqrt{\frac{\text{TP}}{\text{TP+FP}} \cdot \frac{\text{TP}}{\text{TP+FN}}}, \tag{S1}$$

where TP is the number of true positives, FP is the number of false positives, and FN is the number of false negatives.

**Adjusted Rand Index:** Define rand index (RI) as a measure of the percentage of correct decisions made by the clustering algorithm, i.e.,

$$\text{RI} = \frac{\text{TP+TN}}{\text{TP+TN+FP+FN}}, \tag{S2}$$

where TP is the number of true positives, TN is the number of true negatives, FP is the number of false positives, and FN is the number of false negatives.

Then, the Adjusted Rand Index (ARI) can be defined as

$$\text{ARI} = \frac{\text{RI-}E(\text{RI})}{\max(\text{RI})\text{-}E(\text{RI})}. \tag{S3}$$

**Normalized Mutual Information:** Let $U$ and $V$ denote the class labels and cluster labels. Define the entropy of a label set $S$ as

$$H(S) = -\sum_{i=1}^{|S|} P(i)\log(P(i)), \tag{S4}$$

where $P(i) = |S_i|/N$ is the probability of an object in class $S_i$.

The mutual information (MI) between $U$ and $V$ is calculated by:

$$\mathrm{MI}(U,V) = \sum_{i=1}^{|U|}\sum_{j=1}^{|V|} P(i,j)\log(\frac{P(i,j)}{P(i)+P'(j)}), \tag{S5}$$

where $P(i,j) = |U_i \cap V_j|/N$, $P(i) = |U_i|/N$, and $P'(j) = |V_j|/N$.

Then, the Normalized Mutual Information (NMI) is defined as

$$\mathrm{NMI}(U,V) = \frac{2 \times \mathrm{MI}(U,V)}{H(U)+H(V)}. \tag{S6}$$

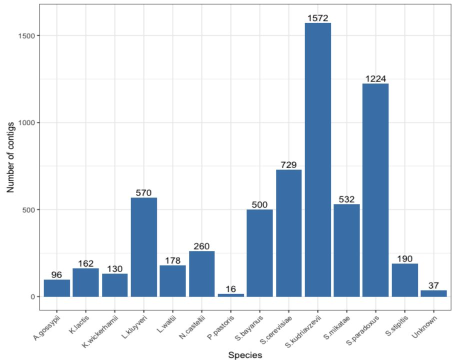


Figure S1: Number of assembled contigs from 13 species.

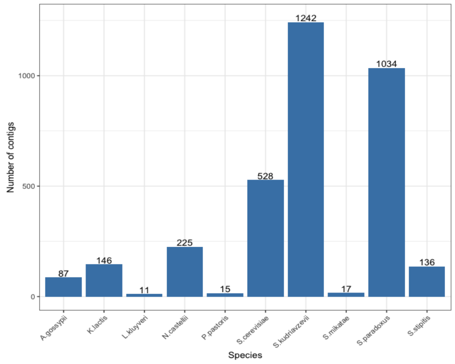


Figure S2: Number of contigs labeled by TAXAssign.

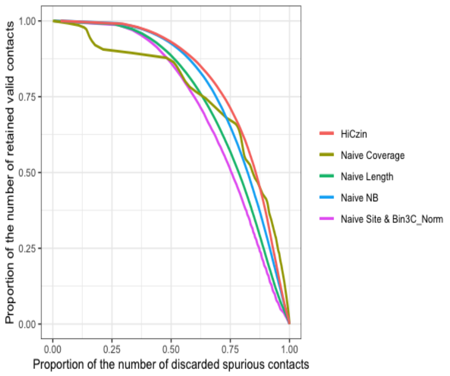


(a)

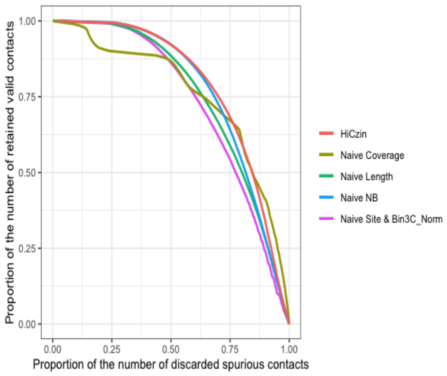


(b)

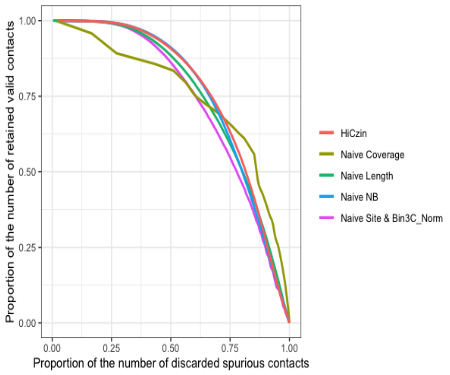


(c)

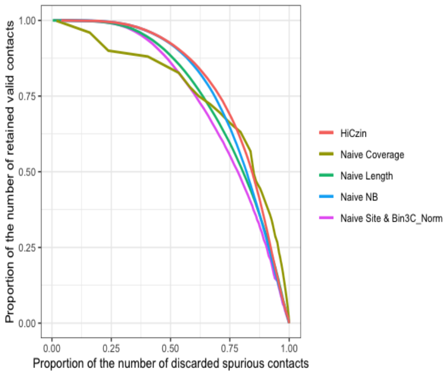


(d)

Figure S3: (a) DR curve using 6 species and 2156 contigs; (b) DR curve using 5 species and 1122 contigs; (c) DR curve using 4 species and 594 contigs; (d) DR curve using 3 species and 369 contigs.

Table S1: M-Y species list in the sample.

| Genus | Species | Strain in sample | Reference strain |
|---|---|---|---|
| Saccharomyces | cerevisiae | FY4H | FY |
| Saccharomyces | cerevisiae | CEN.PK | CEN.PK |
| Saccharomyces | cerevisiae | RM11-1A | RM11-1A |
| Saccharomyces | cerevisiae | SK1 | SK1 |
| Saccharomyces | paradoxus | YDG613 | |
| Saccharomyces | mikatae | FM356 | IFO 1815 |
| Saccharomyces | kudriavzevii | FM527 | IFO 1802 |
| Saccharomyces | bayanus var. uvarum | YZB5-113 | CBS 7001 |
| Naumovozyma | castellii | 4310 | NRRL Y-12630 |
| Lachancea | waltii | Kwaltii ura3 | NRRL Y-8285 |
| Lachancea | kluyveri | FM628 | CBS 3082 |
| Kluyveromyces | lactis | MW98-8C | NRRL Y-1140 |
| Kluyveromyces | wickerhamii | Y-8286 | UCD 54-210 |
| Ashbya | gossypii | WT | ATCC 10895 |
| Scheffersomyces | stipitis | Y-11545 | CBS 6054 |
| Pichia | pastoris | JC308 | GS115 |

Table S2: Number of remaining species, number of remaining contigs and proportion of assigned contigs in each step.

| # of remaining species | # of remaining contigs | Proportion of assigned contigs |
|---|---|---|
| 6 | 2156 | 34% |
| 5 | 1122 | 18% |
| 4 | 594 | 10% |
| 3 | 369 | 6% |

Note: # of remaining species and remaining contigs mean the number of remaining species and remaining contigs.

Table S3: Pearson correlation coefficients(absolute value) between normalized valid contacts and the product of each of the three factors of explicit biases, and AUDRC for different proportions of labeled contigs.

| Proportion of labeled contigs | site | length | coverage | AUDRC |
|---|---|---|---|---|
| 34% | 0.004 | 0.006 | 0.059 | 0.801 |
| 18% | 0.003 | 0.005 | 0.099 | 0.793 |
| 10% | 0.011 | 0.010 | 0.065 | 0.771 |
| 6% | 0.014 | 0.014 | 0.037 | 0.794 |

Table S4: Pearson correlation coefficients(absolute value) between normalized valid contacts and the product of each of the three factors of explicit biases, and AUDRC for different generalized HiCzin models.

| | site | length | coverage | AUDRC |
|---|---|---|---|---|
| HiCzin_LC | 0.006 | 0.002 | 0.097 | 0.812 |
| HiCzin_GC | 0.008 | 0.003 | 0.131 | 0.816 |
| Unlabeled HiCzin | 0.114 | 0.105 | 0.079 | |
